# SIGNAL: Dataset for Semantic and Inferred Grammar Neurological Analysis of Language

**DOI:** 10.1101/2024.11.20.624526

**Authors:** Anna Komissarenko, Ekaterina Voloshina, Anastasia Cheveleva, Ilia Semenkov, Oleg Serikov, Alex Ossadtchi

## Abstract

Recently, the idea of comparison of models’ representations and human brain signals has been a topic of several works. Consequently, several datasets with text data and EEG representations have been published. However, most of the datasets are based on normal reading task with grammatical sentences. At the same time, in the interpretability studies of LLMs, more and more attention is paid to thoroughly designed linguistic tasks based on acceptability measures. In this paper, we present SIGNAL, a dataset for Semantic and Inferred Grammar Neurological Analysis of Language. Our dataset contains a group of sentences with a combination of a fully acceptable sentence and a grammatically or/and semantically incongruent sentences. The dataset has been approved by native speakers and later used for an EEG experiment. In total, our dataset contains recordings of 21 participants, each of whom read 600 sentences. In addition, we present a pilot study where we compare EEG analysis with simple probing experiments.

## Introduction

Aligning LMs and brain language processing has a high potential to be beneficial both for neuroscience and computational linguistics. The question whether a language can be an independent system (1) is still under debate, structured fundamentals of language can and should be discussed in parallel in collaboration of neuroscientists and computational linguists. Current research on brain-model alignment converges on the concept that it is the use of prediction that is shared by both brain and the model and results in partial alignment between the two. A recent study (2) designed a model to predict the extent to which the perceived sentence drive the brain and found that it is not only the predictability (or the associated surprisal) but the subtle aspects related to the form and meaning of the sentence that modulate the speech cortex response strength and therefore highlight specific context in which the brain processes the incoming lexical information.

On the other hand, there is a line of thought in the computational linguistics community regarding the hierarchical predictive modeling implemented in the human language comprehension system (3). At the same time the fact that the language models are effective at predicting brain responses to natural language may not necessarily mean that they perform similar (to the brain) processing but rather capture a wide variety of linguistic aspects (4). Hence, brain-model alignment can reveal how model-based language processing actually resembles human language processing. This can enable answering several valuable questions:

- Whether or not the sentence structure plays a role in generating the surprisal signal?
- Is the contribution of morpho-syntactic parameters to the brain prediction mechanisms similar to that of models?
- Whether processing of semantic and grammatical anomalies are dissociated in humans?

But what data would be useful to compare meaning and grammar? How align the processing taking place in the LLM and that in humans? Predictive coding mechanisms ubiquitously discovered in the nervous system (5) and the corresponding time-resolved neuroimaging studies aimed at exploring brain’s response to the information that is not readily predictable from the context at different processing levels. This make us think that the corpus of semantically and grammatically incongruent sentences with neurolinguistic recordings of humans reading will be instrumental in answering the above questions. The incongruities present in the stimuli will facilitate tracking the processing pathway in humans as the mismatch between the expectation and the measurement modulates brain response to the sensory information. Exploring the layer- and element-wise representations emerging in the LLM and matching them against those measured in humans is likely to highlight similarities and pinpoint the differences in the ways the linguistic information is handled in these systems. The obtained findings could not only be used for improving the LLMs but also will contribute to our understanding of neuronal mechanisms supporting the language function.

Our pilot study provides a valuable tool to do research revealing connections between LMs and neural brain processing. Validation experiments, provided by our study, have already revealed similarities between the two and showed semantic disturbances probably being systematically more notable than syntax in language in general, as evident both on neural level and in LMs activations. We present a carefully created dataset of linguistically acceptable sentences and their semantically, grammatically and both corrupted (incongruent) versions including their normative sources. Our dataset is sourced in 2610 sentences consisting of frequent words sampled from RuSentEval probing suite (6). We simplify these sentences by removing everything but the main clause, and keep resulting sentences of length 3-4 words. We obtain these incongruent counterprats of these sentences in four types of congruity: congruent sentences, semantically incongruent, grammatically incongruent, and semantically-grammatically incongruent. We validate the resulting sentences in a large scale behavioral online experiment to verify the type of stimuli congruity via human assessment. Finally, we choose the top 600 sentences (150 sentences for each congruity type) that have the best online annotators’ agreement over the type of congruity. Having collected EEG data for the chosen sentences, we confirm a significant difference between stimuli congruence conditions on a neuro-physiological level by performing statistical tests on the EEG evoked response data. At the same time, we perform experiments with LLM to estimate layer-wise contrasts between congruence conditions by means of probing. In both cases, we observe semantic errors being seemingly easier to tell than the grammatical ones.

### Related Work

#### Neurolinguistic data in NLP research

The use of neurolinguistic data, such as magnetoencephalography (MEG), electroencephalography (EEG), and functional magnetic resonance imaging (fMRI), can be split in two categories: for using such data in addition to texts to train better models or for studying language acquisition in models and therefore comparing them to human signals.

Hollenstein et al. (7) provide a survey of different NLP applications that involve using brain signals. Hale et al. (8) compare neural models with brain signals to choose a better model. Toneva and Wehbe (9) first probe BERT with brain activity representations and then fine-tune a model with brain data. The more aligned model shows better results than the original model. Similarly, Hollenstein et al. (7) show that combining text data and human signals lead to an improvement on downstream tasks. Ren and Xiong (10) present a CogAlign framework that suggests a new methodology to combine cognitive and text data for better model representations. Murphy et al. (11) use EEG signals to decode part-of-speech tags.

#### Datasets with EEG information

Although there exist a large volume of neuroimaging studies aligning language processing in humans and neural networks, there is still a lack of openly available datasets with carefully created linguistic data balanced on key lexical-semantic properties and high density EEG recordings. The summary comparison of our dataset to existing EEG datasets is presented in Table 2 (Appendix). One of the first datasets was made by Frank et al. (12) to compare language models’ information measures to event-related potentials (ERP) extracted from EEG data. The dataset contains grammatical sentences in English taken from UCL Corpus of reading times (13). ZuCo corpus (14, 15) includes eye-tracking and EEG data taken participants were reading half of the sentences in normal reading set-up and the other half in the task-specific paradigm formulated as a linguistic annotation task. The corpus is made to show differences in human signals between two different setups.

**Table 1.** Key parameters of sentences between groups.

| Sentence structure | Sentence argument | Frequency (IPM), M | Frequency (IPM), SD | Length (syllables), M | Length (syllables), SD |
| --- | --- | --- | --- | --- | --- |
| Subject - Verb - Object | Subject | 3.18 | 0.98 | 102.75 | 135.1 |
| Subject - Verb - Object | Verb | 3.64 | 0.94 | 113.02 | 158.14 |
| Subject - Verb - Object | Object | 3.3 | 1.19 | 120.05 | 167.27 |
| Subject - Verb - Adjective - Object | Subject | 3.2 | 1.0 | 162.41 | 424.47 |
| Subject - Verb - Adjective - Object | Verb | 3.5 | 0.97 | 154.37 | 179.53 |
| Subject - Verb - Adjective - Object | Object | 3.26 | 1.39 | 94.79 | 86.98 |
| Subject - Verb - Adjective - Object | Adjective | 4.0 | 1.13 | 141.28 | 274.98 |
| Subject - Verb - Object - Genitive | Subject | 3.6 | 1.27 | 117.35 | 185.7 |
| Subject - Verb - Object - Genitive | Verb | 3.76 | 0.89 | 123.96 | 200.2 |
| Subject - Verb - Object - Genitive | Object | 3.32 | 1.31 | 129.81 | 207.81 |
| Subject - Verb - Object - Genitive | Genitive | 3.78 | 1.38 | 137.51 | 248.08 |

**Table 2.**
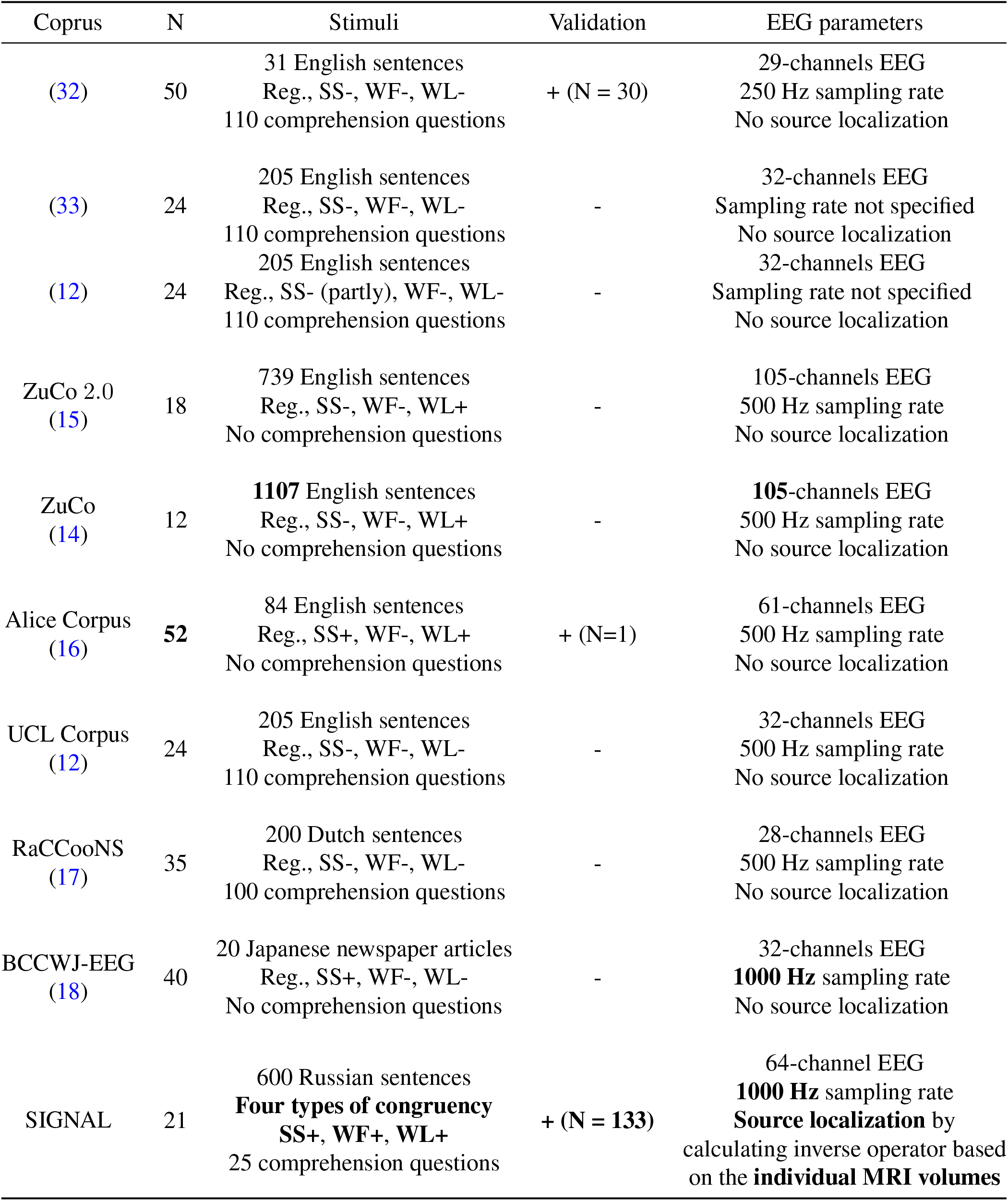
The comparison of our dataset SIGNAL to existing EEG datasets. Reg. refers to regular sentences without incongruent anomalies. SS+/SS-refers to controlled/not controlled syntactic structure. WF+/WF-refers to controlled/not controlled word frequency in the sentences. WL+/WL-refers to controlled/not controlled word length in the sentences. If the corpus was validated, N refers to the number of participants in the validation study.

| Coprus | N | Stimuli | Validation | EEG parameters |
| --- | --- | --- | --- | --- |
| (32) | 50 | 31 English sentences<br>Reg., SS-, WF-, WL-<br>110 comprehension questions | + (N = 30) | 29-channels EEG<br>250 Hz sampling rate<br>No source localization |
| (33) | 24 | 205 English sentences<br>Reg., SS-, WF-, WL-<br>110 comprehension questions | - | 32-channels EEG<br>Sampling rate not specified<br>No source localization |
| (12) | 24 | 205 English sentences<br>Reg., SS- (partly), WF-, WL-<br>110 comprehension questions | - | 32-channels EEG<br>Sampling rate not specified<br>No source localization |
| ZuCo 2.0<br>(15) | 18 | 739 English sentences<br>Reg., SS-, WF-, WL+<br>No comprehension questions | - | 105-channels EEG<br>500 Hz sampling rate<br>No source localization |
| ZuCo<br>(14) | 12 | **1107** English sentences<br>Reg., SS-, WF-, WL+<br>No comprehension questions | - | **105**-channels EEG<br>500 Hz sampling rate<br>No source localization |
| Alice Corpus<br>(16) | **52** | 84 English sentences<br>Reg., SS+, WF-, WL+<br>No comprehension questions | + (N=1) | 61-channels EEG<br>500 Hz sampling rate<br>No source localization |
| UCL Corpus<br>(12) | 24 | 205 English sentences<br>Reg., SS-, WF-, WL-<br>110 comprehension questions | - | 32-channels EEG<br>500 Hz sampling rate<br>No source localization |
| RaCCooNS<br>(17) | 35 | 200 Dutch sentences<br>Reg., SS-, WF-, WL-<br>100 comprehension questions | - | 28-channels EEG<br>500 Hz sampling rate<br>No source localization |
| BCCWJ-EEG<br>(18) | 40 | 20 Japanese newspaper articles<br>Reg., SS+, WF-, WL-<br>No comprehension questions | - | 32-channels EEG<br>**1000 Hz** sampling rate<br>No source localization |
| SIGNAL | 21 | 600 Russian sentences<br>**Four types of congruency**<br>**SS+, WF+, WL+**<br>25 comprehension questions | + (N = 133) | 64-channel EEG<br>**1000 Hz** sampling rate<br>**Source localization** by<br>calculating inverse operator based<br>on the **individual MRI volumes** |

Unlike previous dataset, Alice Corpus (16) contains data about fMRI and EEG signals while participants were reading the first chapter of *Alice in Wonderland*. Therefore, the dataset allows to compare human signals to computational models on a text with narrative structures.

As for other languages, Frank and Aumeistere (17) present a dataset of narrative sentences in Dutch and Oseki and Asahara (18) created a corpus of newspaper articles with human signals for Japanese.

Still, the majority of previous datasets were recorded based on English normal sentences, and there were no other datasets exploring processing of anomalous language data. On the other hand, Russian language is flective and characterized with a wide range of morphological categories (such as tense, gender, number, person, case) and a complicated inflection system as opposed to analytical languages that use specific grammatical words rather than inflection (for example, English). In the current study, we provide linguistic material of processing of both semantical and grammatical anomalities, and the regularities discovered using our dataset are likely to be extendable to many other flective languages like Slavic, Romanian, German etc. Secondly, the majority of previous studies did not control stimuli for key lexical-semantic and morphosyntactic properties such as syntactic sentence structure, word frequency and length. On the other hand, the quality of stimuli is crucial for neurophysiological experiments. Sentence structure significantly influences brain responses (19–21) and processing of particular words can be even supported by different neural networks depending on the part of speech and argument structure (22). In the current study we provide providing a dataset with well-controlled stimuli balanced on key lexical-semantic properties and controlled syntactic structure, which can enable clearer insights into the differences observed in response to various morpho-syntactic properties, enhancing the reliability and interpretability of both LMs probing and neurophysiological EEG results.

### Corpus collection

#### Data generation

Our data is based on RuSentEval dataset (6), the probing suite for Russian language. For each sentence, we marked up a sentence structure using Natasha Python library and reduced^1^ each sentence to one of three structures if possible: subject + verb + object, subject + verb + adjective + object, or subject + verb + object + genitive. For each word in the sentence we extracted frequency based on Russian National Corpus^2^ and length in phonemes and syllables.

To obtain sentences of four congruency types, we generated three variants of each sentence with semantical, grammatical, and semantical-grammatical errors by replacing the Object in each sentence. Firstly, we generated semantically incongruent sentences by replacing a target word (Object within the sentence structure) in a congruent sentence with another word. We take `word2vec` to generate negative examples, i.e. with words furthest from the original one in a vector space. As for implementation, we use the model from RusVectores package^3^ (see Algorithm 1). Then we took congruent and semantically incongruent sentences to generate sentences of other two types: with a grammatical error and with both types of errors respectively. We intentionally put the object of each sentence in the wrong case or number (see Algorithm 2). To inflect the wordform acaccordingly we use the pymorphy2^4^ tool.

In the result, we obtained sets of sentences that differ from each other only by congruency type of Object argument (precisely, presence or absence of a grammatical/semantical error). For example:

- **Congruent sentence**: *Storony podpisali <u>soglashenie</u>*. ‘The parties signed <u>an agreement</u> (accusative)’
- **Semantically incongruent sentence**: *Storony podpisali <u>detstvo</u>*. ‘The parties signed <u>childhood</u> (accusative)’
- **Gramatically incongruent sentence**: *Storony podpisali soglashenii*. ‘The parties signed an agreement (locative)’
- **Semantically and gramatically incongruent sentence**: *Storony podpisali <u>detstve</u>*. ‘The parties signed <u>childhood</u> (locative)’

In total, we obtained 10440 sentences stimuli of four types of conditions for further evaluation. Based on the results of online validation, we choose 600 sentences (150 congruent ones with three incongruent variants). Congruent sentences were divided into three syntactic structures (50 “congruent” sentences in each group); sentence arguments balanced in terms of frequency and length between groups (i.e, verbs are balanced with verbs, subjects with subjects, objects with objects between groups). Thus, the corpus contained 4 main (congruency) categories with 3 (syntactic structure) subcategories each resulting in a total of 12 well-balanced subcategories.

#### Human evaluation

##### Annotation setup

To collect humans’ acceptability judgements for the generated sentences, we use a crowdsourcing platform Toloka^5^. The task of crowd workers is to evaluate the quality of data generation. The workers were asked to assess if a given sentence is congruent, it contains either a semantic or grammatical error, or it contains both types of error (for the example of web interface see Appendix and for the instructions given to crowd workers see Appendix). The task was only available for people who listed Russian as their native language and come from Russia, Belarus, Ukraine or Kazakhstan (based on their phone number).

##### Quality control

As for the quality control, we use control tasks. Both training and control tasks were annotated by the authors. We use the following constraints for workers to ensure faithful annotation:

- they are only allowed for a task if they annotate the held-out 26 sentences with at least 90% accuracy
- if they earned 5 dollars or more, are suspended from the task for 2 days (as they completed too many tasks);
- if they skipped 7 tasks or more, they are banned for 2 hours;
- if the quality on control tasks is less than 50% out of 4 recent responses, they are banned for 2 days;
- if 30% of responses are different from other annotators’ responses, they are banned for 3 days.

##### Annotation statistics

Each task was annotated independently by 3 crowd workers. The overlap increased if any of crowd workers was banned.

In total, 133 people annotated sentences with average of 6.95 per person. The price for each task was $0.014. The overall number of annotated sentences^6^ was 1465 that were later filtered out if less than 75% of crowd workers noticed an error. For the further experiments, we form sets of 4 sentences, where one of the sentences was acceptable, one is a modified version of the same sentence with a semantic error, one version contains a grammatical error and one version has both types of errors. We also balanced out the dataset by sentences syntactic structure. After filtration, we have 150 such sets of sentences, where 50 sets have the structure subject + verb + object, 50 sets have the structure subject + verb + adjective + object, and 50 sets follow the structure subject + verb + object + genitive.

### EEG data collection

#### Participants

Data of 21 participants (*M*_*age*_ = 21.76, *SD* = 3.66, female = 16) were collected for the corpus. All participants were paid a standard rate $2.7 per hour. All participants were right-handed native Russian speakers, did not have a linguistic background and were not proficient speakers of any foreign language other than English. They had normal or corrected vision and hearing, no history of neurological deficits or language-related impairments. Participants signed a written informed consent form prior to taking part in the research and completed the screening questionnaire form to estimate language proficiency and reading skills and reveal any contraindications. The study was approved by the local university Committee on Interuniversity Surveys and Ethical Assessment of Empirical Research.

According to the screening questionnaire, all participants had high proficiency in Russian, as they had good or excellent grades in school in Russian language and all participants were well-read. All participants had at least 11 years of education, which is equal to upper secondary education (*M*_*age*_ = 14.26 years of education, *SD* = 1.74).

#### Procedure

Experimental task consisted of 600 sentences of four congruence conditions (150 sentences per condition) each divided into three subcategories according to sentence syntactic structure. Additionally, 25 control sentences with a context-related questions were included into the task to control for participants’ attention.

Experiment started with a participant preparation and electrode montage. Participants were instructed to read sentences attentively and to answer to sentence-related questions after a part of the sentences. The task was presented on the screen in the beginning of the experiment for a second time. During the experiment, each sentence started with a fixation cross lasting for 0.5 seconds. Then sentences appeared on the screen word by word with a speed one word per 0.5 seconds close to natural reading speed (23). The order of congruency type and positions of fillers in relation to the target sentences were randomised in order to prevent participants from getting a clue of the experiment. Counterparts of similar sentences (differing only by congruency type) were set on a maximum distance so there were at least 100 other sentences between two similar items. There were six 30-second breaks during experiment after every 75 sentences (every 2-3 minutes) and a long 3-minute break in the middle of experiment to check the impedance of the electrodes and give participants time to rest and refresh their attention. The total duration of experiment was 2 hours including preparation time. The outline of experiment is visualized in Figure 1.

**Fig. 1.**
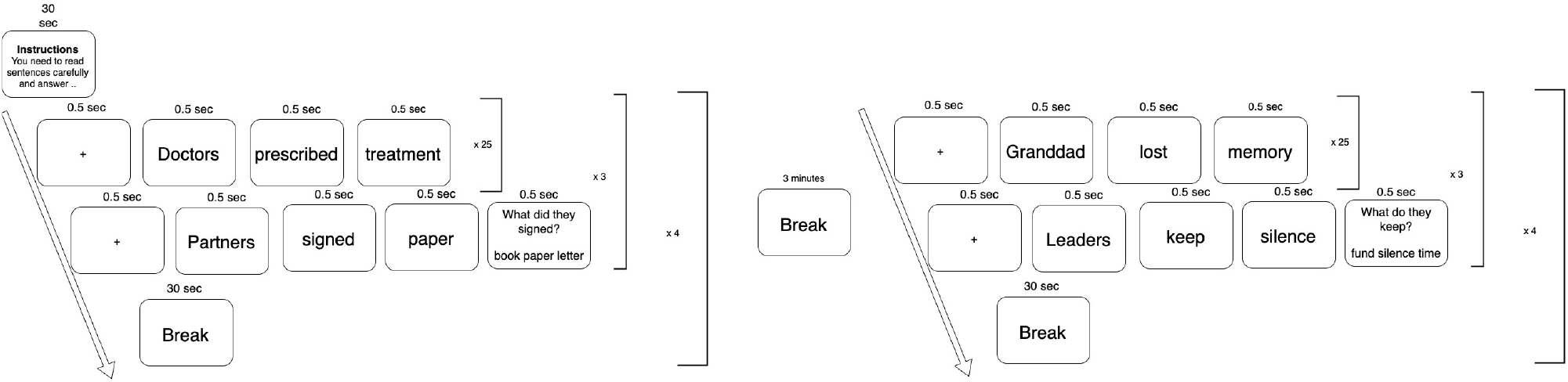
The outline of the experiment.

#### EEG recordings

Participants were fit with a 64-electrode cap (AntiCap, BrainProducts, German) in a sound attenuating, electrically shielded booth with a monitor and a keyboard. Electrodes were positioned on the scalp according to the International 10–20 system with a Cz electrode used as a reference. The task was programmed and presented via PsychoPy (v2023.2.2) (24). EEG data were collected using an ActiCap electrode system (BrainProducts, Germany) and actiCHamp amplifier (BrainProducts, Germany) at 1000 Hz sampling rate with impedance below 10 kΩ.

#### EEG preprocessing and analysis

EEG data preprocessing was performed via MNE Python library (25). A low-pass zero-phase filter with a 30-Hz cutoff, and a high-pass filter with a 0.1-Hz were applied to the EEG recordings. Based on their spectral characteristics, noisy/flat channels were interpolated. Independent component analysis (ICA) was performed to remove ocular artifacts such as horizontal and vertical eye-movement. The data were divided into epochs containing only the target stimuli recordings, including 400 ms pre-stimulus and 800 ms post-stimulus interval. The raw EEG data include mark-up with a unique ID for each sentence and a congruency condition of the target word. Baseline correction was applied to the epochs using 0-th order detrend computed over the prestimulus (−400 ms - 0 ms) interval.

Statistical analysis was aimed to reveal statistically different temporal-spatial clusters of EEG data between four stimuli congruency conditions. To do so, we firstly obtained event-related potentials (ERP) (26) data by averaging EEG response between participants and conditions. Secondly, computed mean values and standard deviations of ERP and estimated z-scores to assess the presence of differences between each pair of condition. To estimate a statistical significance of results, a simple non-cluster based permutation test was performed on the ERP maps. Grand-average data were obtained via averaging z-scores between participants, p-values values were combined with a Fisher’s combined probability test. Benjamini-Hochberg (27) procedure was applied to correct for multiple comparisons using the false discovery rate (FDR) correction procedure with the critical alpha level set to 0.016.

### Data validation results

The statistical analysis was aimed to ensure our dataset satisfies the necessary conditions for that kind of data. Firstly, we estimated EEG data to reveal distinguishable difference between stimuli congruence types on neurophysiological level. Secondly, we estimated whether LLMs intermediate activations allow to distinguish these groups of sentences. As our human recordings were made on carefully curated *subset* of the larger dataset derived from RuSentEval, we experiment with LLMs on both initial dataset and the curated subset.

#### EEG recordings validation

Participants’ performance in a behavioral control task was high (*M* = 0.81, *SD* = 0.20). The results of z-statistics reveal significant difference in all pairs of conditions. Figure 2 represents a topography of relevant statistically significant spatial-temporal clusters based on the permutation tests corrected for a false-discovery rate. The results of z-statistics reveal significant difference in all pairs of conditions. Figure 2 represents a topography of statistically significant spatial-temporal clusters based on the permutation tests corrected for a false-discovery rate. According to 5, the grammatically incongruent sentences cause strong negative (P in the EEG terminology) responses in the interval around 400-600 ms post stimulus. Semantically incongruent sentences have a strong positive response (N in the EEG terminology) in this time interval observed in (largely) the complementary set of channels. At the same time, it seems that the positive (N) response in the semantically incongruent condition is of shorter duration and may still represent N400 component, whereas longer lasting negative deflection in the response to the grammatically incorrect sentences may be the expected P600 component.

**Fig. 2.**
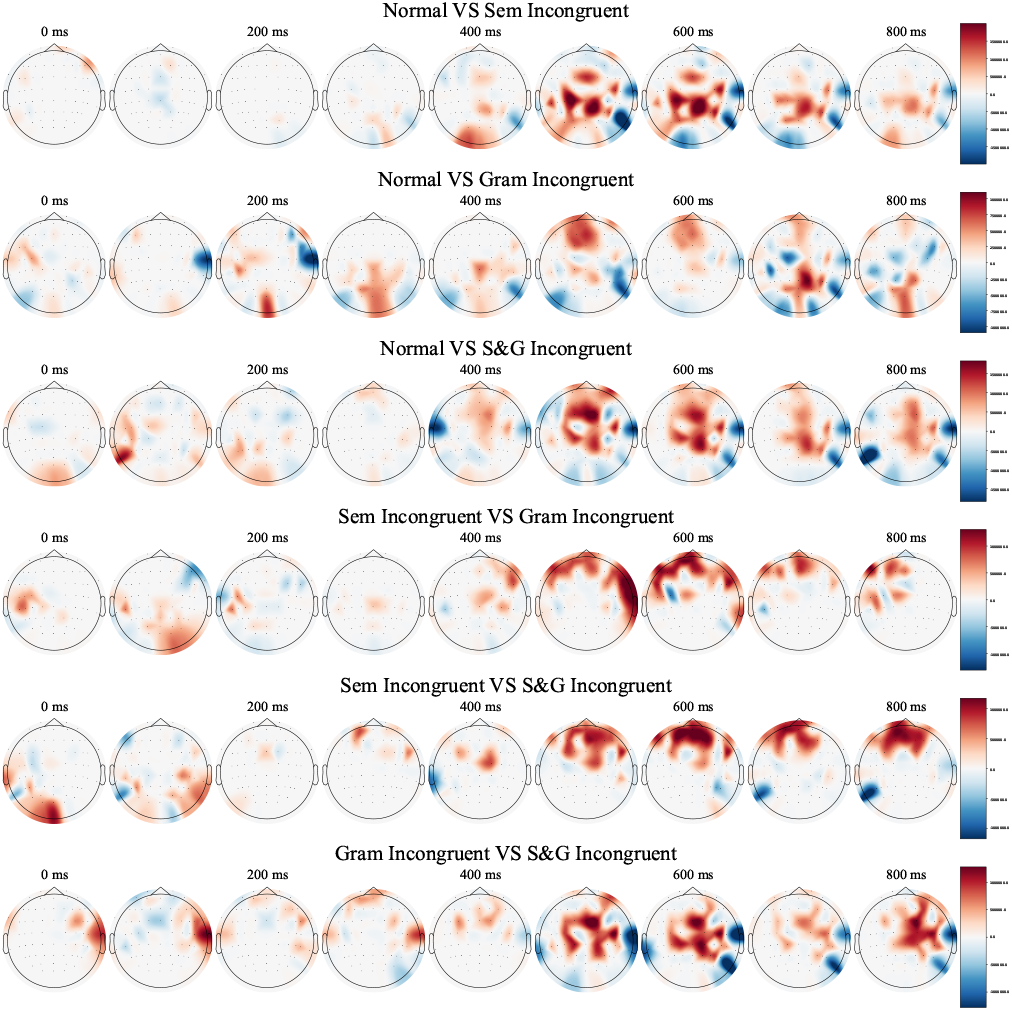
Topographical plots of z-scores of significantly different ERP between conditions presented over a 100 ms time window

#### Corpus validation with LLMs probing

To perform the validation of our corpus with probing experiments, we collect activations of each layer of the ruBERT LLM (28), a de-facto standard LLM for processing Russian texts. The model follows the standard BERT architecture of 12 layers, each consisting of 768 neurons. Probing experiments were performed using the Google Colaboratory free compute service and took approximately one hour of computation in total.

These experiments aim to verify whether the LLM in fact exhibits differential activation in response to different types of errors in the dataset. This step confirms the dataset’s suitability for aligning LLM representations with EEG data, as further analysis would be irrelevant without observed differentiation. The experimental design follows the generic probing setup as in work of e.g. Conneau and Kiela (29). We perform layer-wise probing, with logistic regression probes employed to tell the congruity type of a sentence. We mitigate the effect of random seeds in our small-scale experiments by performing each experiment 20 times. As sentence representations, we use contextualized vectors of the word bearing the congruity/incongruity of a sentence. If a word consists of several subtokens, the respective embeddings are averaged. See more details on tokenization safety-check below.

##### Tokenization study

When creating incongruent sentences, most of the time we did not introduce more tokens nor reduce the number of subwords according to the tokenizer used in the model. Only in 7.5% of all cases, semantically incongruent sentences were one token longer than the original ones. Similarly only in 8.5% of all cases, the same thing is observed for grammatically incongruent sentences. Thus, there is no way for a probing model to tell the incongruence by revealing the length of a sentence in tokens.

##### Experiments results

The aim of telling one congruity type from another for a sentence leaves us with six binary pairwise classification probing experiments ^7^, results of experiments are shown in Figure 3. We find middle-to-high layers’ activations to be most prominent to detect congruity types in a sentence. Semantically (Sem) incongruent sentences are easier to detect than grammatically (Gram) incongruent ones, at the same time Semantically *and* grammatically (S&G) incongruent sentences are the easiest to tell from normal ones, and are the hardest to tell from both Sem and Gram incongruent sentences.

**Fig. 3.**
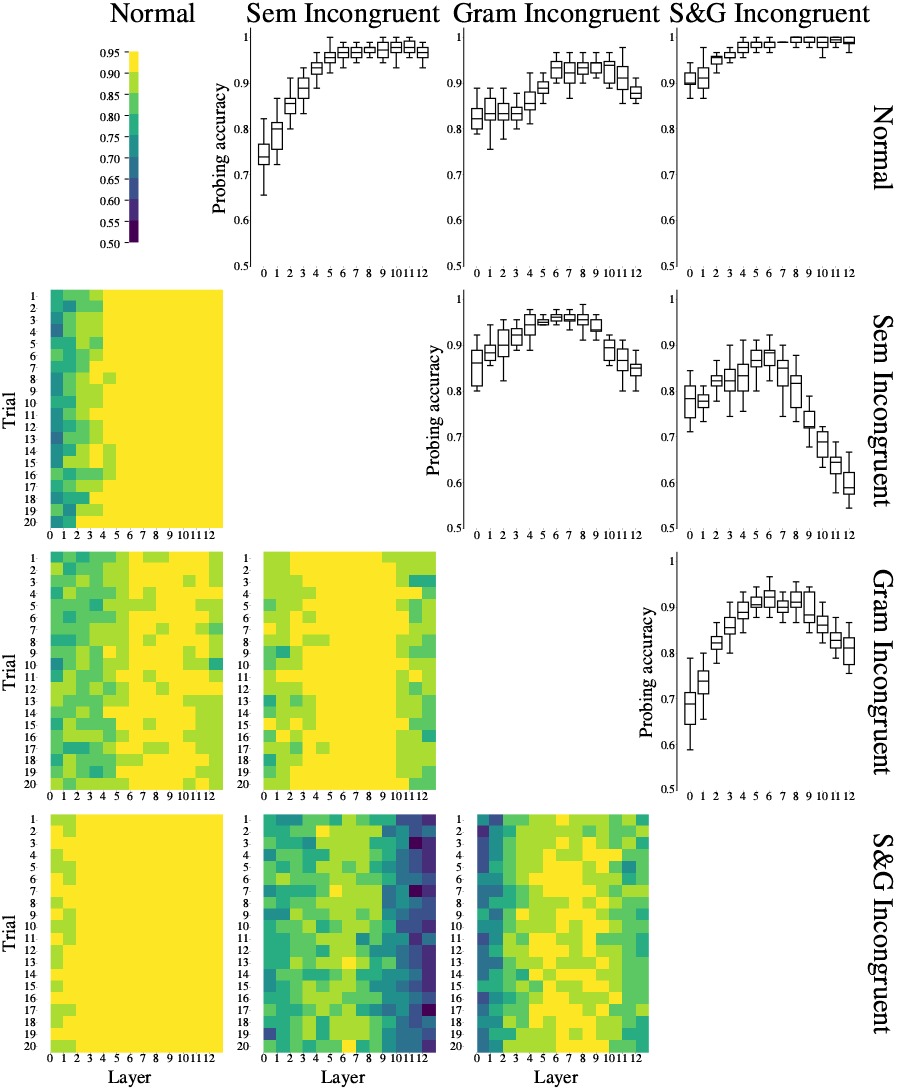
Layerwise probing scores for the incongruence detection probing task. Semantically incongruent sentences thus appear more “weird” to the model than the grammaticaly incongruent ones. The case of dual incongruence is even easier to tell.

While probing experiments so far were performed with our proposed SIGNAL dataset, we also perform the same experimental pipeline with the sentences initially sampled from RuSentEval and their incongruent counterparts. Results on this larger dataset support our findings on SIGNAL dataset. In particular, once again we find semantically incongruent sentences appear more surprising for the model than the grammaticaly incongruent ones. The case of dual incongruence is even easier to tell. Results charts can be found in appendix. We applied Representational Similarity Analysis (30) to evaluate activation difference between 12 types of stimuli (three groups of sentences different by syntax structure each divided into four congruency conditions) detected by LLMs. As a result, we obtained layerwise Representational Dissimilarity Matrices (RDMs) contrasting each pair of condition presented in Figure 4. As can be observed from the RDMs, they demonstrate different kinds of representations. The RDM from the first layer reflects congruency types while the RDM from the one before the last 12-th layer elicits and sentence structure, see section.

**Fig. 4.**
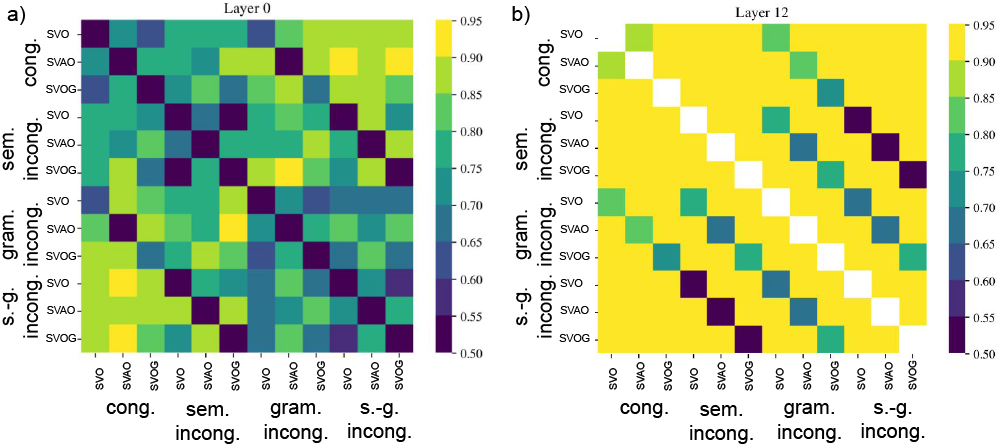
RDM from the 1st a) and the 12th b) layer of the LLM derived from the probing experiment. Higher classification accuracy for each pair of conditions corresponds to the larger distance between the layer activation profiles.

**Fig. 5.**
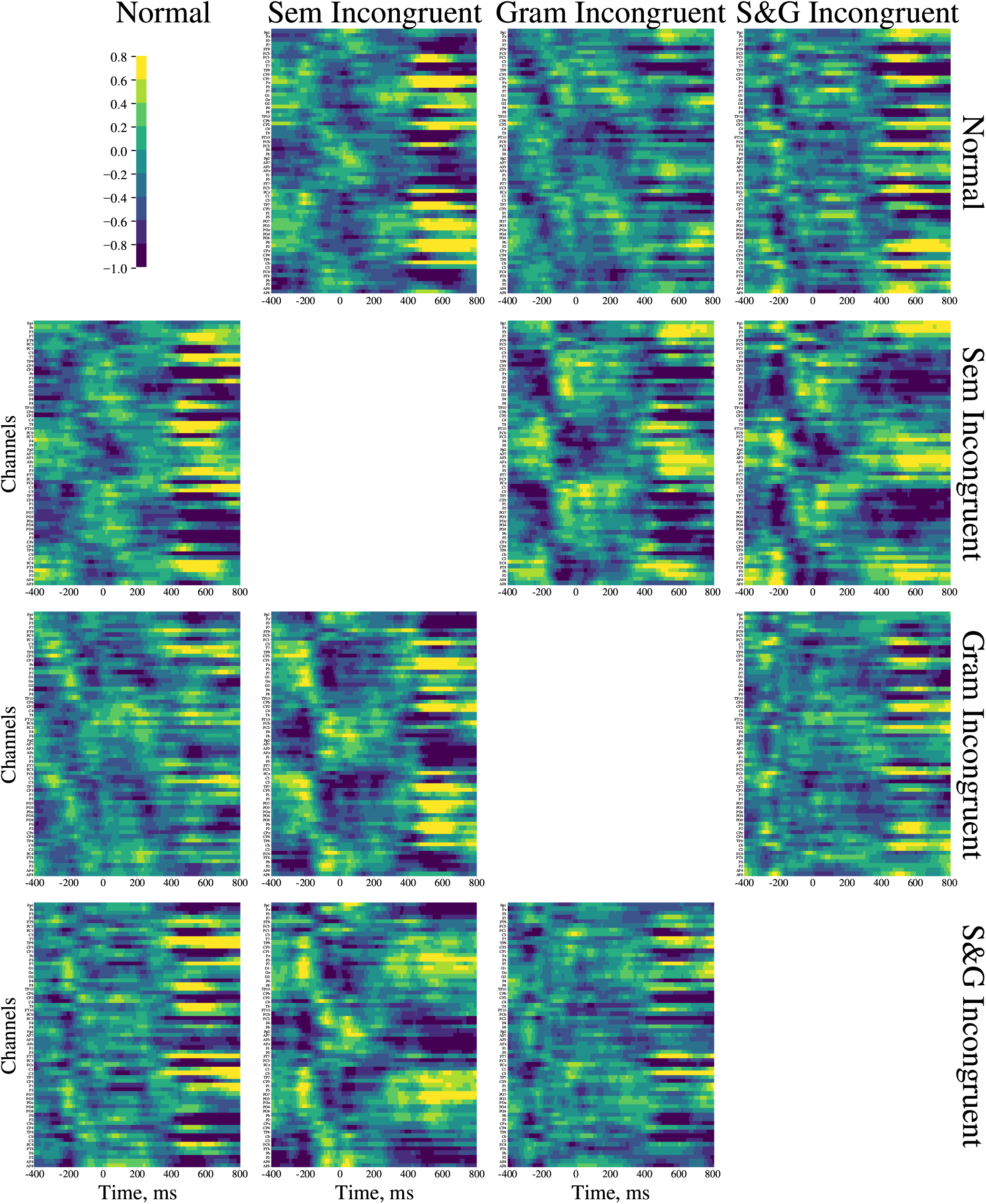
Results of pairwise z-scores for EEG recordings. The statistically significantly differing intervals are depicted at the Figure **??**

**Fig. 6.**
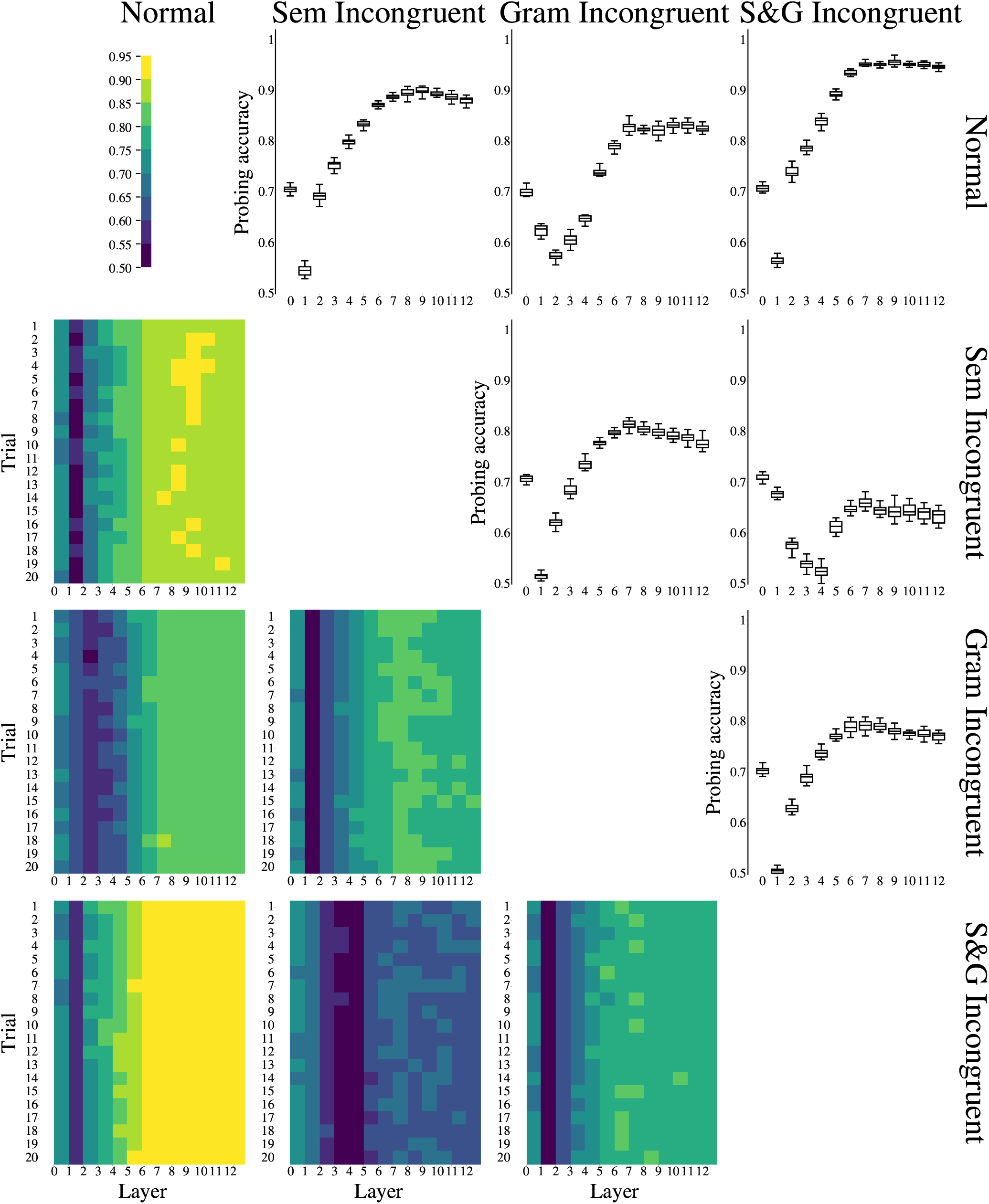
Results of layerwise probing for the task of pairwise congruity types detection based on layer activations. Lower triangle: results over each of the 20 independent runs made to mitigate the effect of random seed. Upper right triangle: boxplot representation of the same data, aggregating over independent run results.

## Discussion

The current study presents dataset which is primarily valuable computational linguistics by its quality and well-balanced stimuli parameters as a base for an ultimate goal of brain-model alignment. The preliminary tests demonstrated the presence of significant topically organized neurolinguistically plausible differences in the EEG data between incongruity conditions. We have also probed an LLM with the developed corpus of sentences. Based on our observations, the tested LLM clearly detects the presence of incongruency combined with the observed structure assuming the data validity for further investigations.

Firstly, the current dataset is unique as it consists of well-balanced both regular and anomalous sentences with grammatical and semantic errors, see Table 2 (Appendix) for comparison with the existing datasets. The stimulus includes well-structured and verified language material including sentence groups distinguished by three syntactic structures and four congruency conditions (semantical, grammatical, and semantical-grammatical). Anomalous stimuli were generated using language model, and validity of them was checked via a web-based validation study with 133 respondents to prove that congruence/incongruence type is correctly identified by Russian native speakers.

Assuming the well-balanced structure of the dataset, it can be easily divided into four main categories with three subcategories each resulting in a total of 12 well-balanced subcategories. This prospectively enables the RSA (Representational Similarity Analysis) including its recent modifications permitting a computationally light analysis (31) to bridge the gap between language models and the cortical processes.

Finally, the neuro-physiological data in the current dataset was collected by means of high density EEG. These high quality measurements of brain activity combined with a well designed set of stimuli and individual MRI scans positively distinguishes this dataset from the existing ones and allows for a sufficiently spatially-detailed and time-resolved modeling of neuronal sources to track the dynamics of the prediction errors and matching those against the processes unfolding in a language model. This will help to resolve the reasons behind language models being effective at predicting brain responses to natural language and design novel architectures and training policies to better align artificial agents with human beings.

## Conclusion

We have presented the first dataset comprising carefully selected and crowd-verified short sentences with varying kinds of incongruity along with the 64-channel EEG recordings from humans reading these sentences in a carefully designed experimental paradigm. The presented dataset and the accompanying tools will facilitate research into the neural bases of language processing in humans in alignment with NLP performed in LLMs.

## Limitations

Our dataset contains information that is limited to a specific group of people (see Section) and might be different from results of experiments with different target groups. Moreover, the dataset might include topic biases which may have affected the results. Another limitation is that the EEG data are collected from only 21 subjects which is currently being ameliorated by recording more volunteers to ensure the gender balance and reduce possible other biases. Yet, given the extent to which the parameters of this dataset are controlled it represents the most comprehensive resource available today designed for the purposes explained above.

## Ethical considerations

All participants of EEG experiment signed an informed consent form and were made aware of purposes of the experiment. The study was approved by the ethical committee of our institution. We do not publish any personal data in our dataset.

## Comparison to existing EEG datasets

### Crowd-sourcing task interface

#### Annotation interface in Toloka

The example of annotation task is the following.

Target sentence: *Economists make positive forecasts\** (the word *forecasts* here is in a wrong case)

Translation of the question: *Is there an error in this sentence and what kind of error is it?*

Translation of the answer options:

1. *No, there is no error*.
2. *Yes, there is an error in this sentence. The sentence is meaningless*.
3. *Yes, there is a grammatical error in this sentence*.
4. *Yes, there is a grammatical error in this sentence and it is meaningless*.
5. *Yes, there is an error in this sentence but it is hard to specify the error’s type*.

#### Algorithms for generation of semantically and grammatically incongruent sentences

##### Algorithm 1

Generation of semantically incorrect sentences

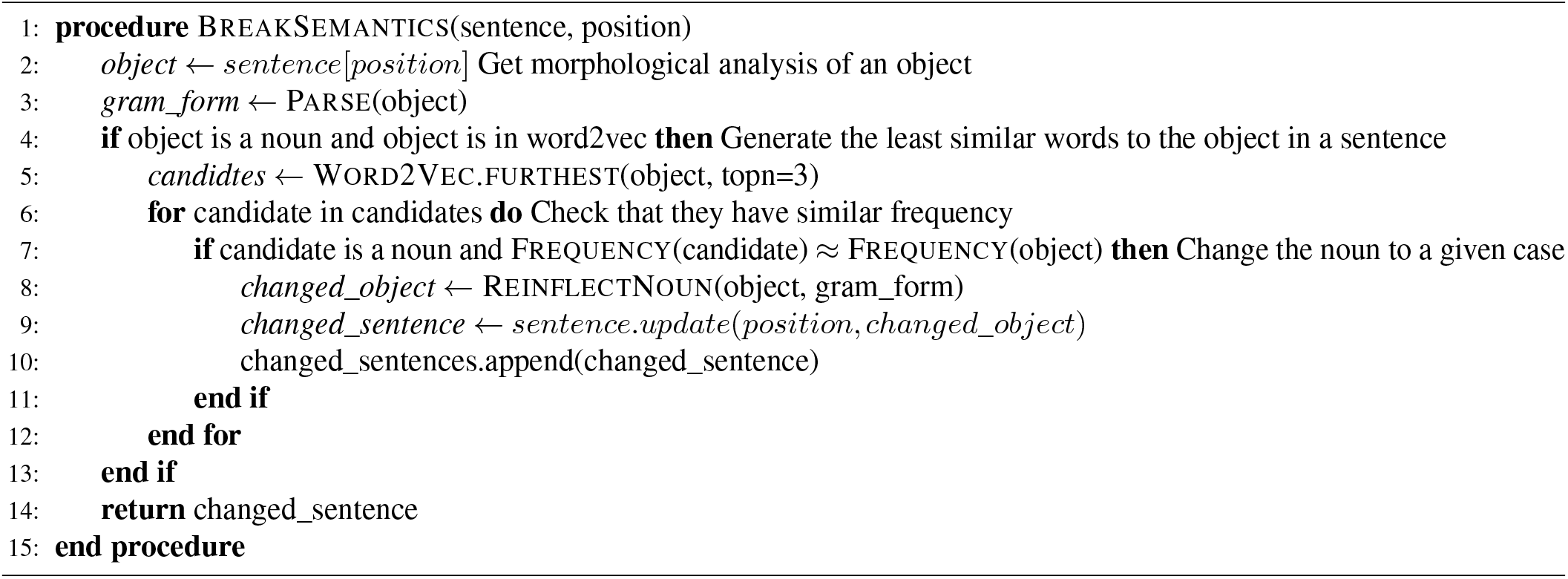

##### Algorithm 2

Generation of grammatically incorrect sentences

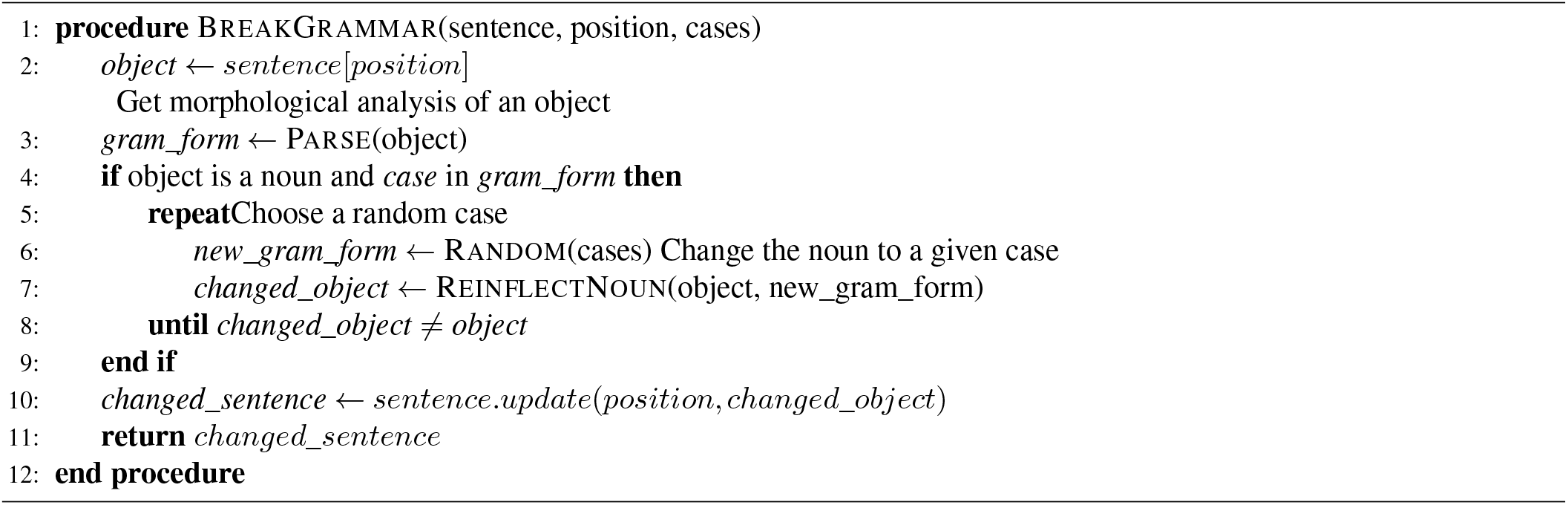

#### Instructions for crowd workers

Read the sentence. Assess if there is an error in this sentence. If there is one, specify its type.

##### Types of errors

- **The sentence is meaningless**: choose this type if you think that the sentences is meaningless, for example:
  - *Avtobusy prohodjat massovuju fortunu*. Buses are subject to massive fortune.
- **The sentence contains a grammatical error**: choose this option if one of the words in the sentence is in a wrong case or number, for examples:
  - *Numeracija chashhe vsego proishodit natural’nymi chislom*. Numbering most often occurs with natural number.
  In this sentence a word is used in a wrong number, the correct way would be to use the word *chislami* ‘numbers’.
  - *Posmotrite na nashi roditeli*. Look at our parents.
  In this example, the word is in the wrong case, the correct way of saying would be *na nashih roditelej* ‘at our parents’.
- **Hard to specify the error’s type**: if the sentence seems incorrect to you but the error is different from the errors described above, choose this option. Please choose this option only in the case if you are certain that this error can be classified with categories mentioned above.

Notice that a sentence can be fully acceptable, in this case you do not have to specify any type of an error. Moreover, sentences could contain both types of the errors:

- *Glavnyj geroj otpravljaetsja v trude*. The main character is going to the hard work.

In this example, the word *hard work* does not fit into the sentence based on its meaning (it is not possible to go to *hard work*) and this word is in a wrong case (it should be accusative).

#### Data storage

https://huggingface.co/datasets/zhuravlevahana/SIGNAL

#### Analytics and visualizations code

https://github.com/AIRI-Institute/SIGNAL

#### Pairwise z-scores between conditions for the spatio-temporal EEG recordings

## Footnotes

1 In supplementary experiments (Section) we show that dataset reduction did not spoil the observed probing results on the dataset, thus, preserving the general applicability of the data for probing.

2 https://ruscorpora.ru/en/

3 https://rusvectores.org/en/models/

4 https://pymorphy2.readthedocs.io/en/stable/index.html

5 https://toloka.ai/

6 Given economic constraints and initial goal to get 150 sentences per condition, we did not annotate the whole machine-generated set.

7 congruence types are described in section

